# Theory of microtubule length regulation in antiparallel overlaps

**DOI:** 10.1101/743880

**Authors:** Hui-Shun Kuan, Meredith D. Betterton

## Abstract

During cell division, microtubules in the mitotic spindle form antiparallel overlaps near the center of the spindle. Kinesin motor proteins alter microtubule polymerization dynamics to regulate the length of these overlaps to maintain spindle integrity. Length regulation of antiparallel overlaps has been reconstituted with purified microtubules, crosslinkers, and motors. Here we develop a theory of steady-state overlap length which depends on the filament plus-end motor concentration, determined by a balance between motor arrival (motor binding and stepping in the overlap) and motor departure (motor unbinding from filament tips during depolymerization) in the absence of motor-driven sliding. Assuming that motors processively depolymerize and exhibit altered binding kinetics near MT plus-ends improves the agreement between theory and experiment. Our theory explains the origin of the experimentally observed critical concentration, a minimum motor concentration to observe a steady-state overlap length.

---

To regulate the size of organisms, tissues, cells, and subcellular structures, biological organisms must solve fundamental physics problems of measuring and controlling their size. Cells use molecules that are nanometers in size to sense and regulate lengths of microns to tens of microns. One elegant method to bridge this separation of scales uses cytoskeletal filaments, which are structurally polar, micron-scale quasi-one-dimensional polymers tunably constructed from nanometer-scale building blocks. Length regulation of cytoskeletal filaments is biologically important for determining organelle and cell size [1–4]. Motor proteins can regulate filament length by binding to filaments, walking toward one end, and altering filament polymerization dynamics at the tip: because accumulation of motors at the tip can depend on filament length, this mechanism allows length sensing and regulation [5–11].

In addition to length regulation of single filaments, overlaps between two filaments are regulated in length. For example, the mitotic spindle that segregates chromosomes during eukaryotic cell division is formed from polymers called microtubules (MTs) in a bipolar structure, with MTs from each half of the spindle meeting in its center and forming stable antiparallel overlaps (fig. 1A). These overlaps are important for spindle force generation and are regulated in length [12–15]. Previous work has examined how antiparallel sliding motors such as kinesin-5 motors, which tend to separate overlapping filaments, and crosslinking proteins from the PRC1/Ase1/MAP65 family, which tend to increase overlap length [16], can balance to give stable overlaps [17–19]. Length-regulated overlaps can be created in vitro via crosslinkers and motors that bind to the crosslinkers and exert force on them [20–22]. In addition, Bieling and Surrey experimentally demonstrated regulated antiparallel MT overlaps in a system in which little to no sliding occurs [23]. In this system, length regulation appears to occur analogously to the single-filament case by kinesin motor proteins that are recruited to the overlap by crosslinkers, then walk to MT plus-ends and slow their growth [23]. Therefore, minimal ingredients of growing MTs, crosslinking proteins, and motors can produce antiparallel overlaps of stable length. However, the precise mechanisms by which motor activity at the MT plus ends leads to stable overlap length are not clear.

**FIG. 1.**
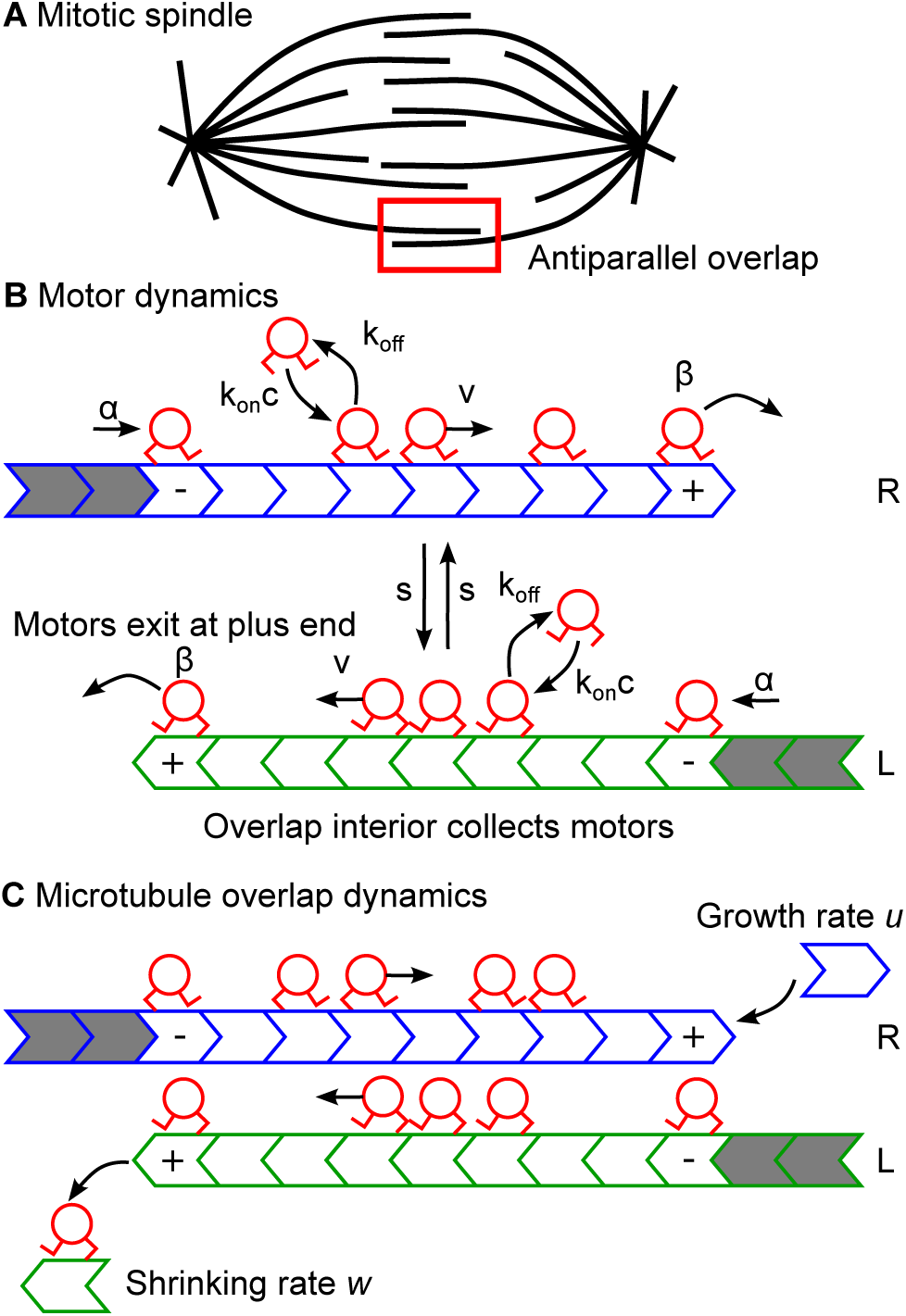
Spindle and model schematic. (A) Mitotic spindle schematic, with microtubule antiparallel overlap region labeled by red box. (B,C) Model schematic, with overlap sites in white, MT sites outside overlap in grey. (B) Motor dynamics, including binding/unbinding, plus-end-directed stepping, and switching in the bulk. At boundary sites, overlap entry (minus-end) and exit (plus-end) occur. (C) Microtubule dynamics in the overlap. Growth and motor-dependent shrinking occur. R/L label right/left direction of motor motion; +/- label MT plus=/minus-ends.

Theoretical models of motor protein traffic and length regulation have built on the totally asymmetric simple exclusion process (TASEP) [24–27], and have successfully predicted features of motor density profiles on MTs [28, 29]. For motors moving left to right on a single filament, the density profile depends on the boundary conditions, leading to three phases: the low-density phase (left-boundary dependence, abbreviated IN), the high-density phase (right-boundary dependence, abbreviated EX), and the maximum current phase (abbreviated MC) [24, 25, 30]. If the length of the filament is dynamic, MT polymerization creates a different boundary condition which drives the overall density profile to one of these three phases, and might make the steady-state length unstable [10, 11, 31–33]. Therefore, it is interesting to examine the physical properties that make overlap length stable or unstable.

Here we develop a theory of overlap length regulation induced by motor proteins that shorten MTs at their plus-ends, inspired by Bieling and Surrey’s experiments [23]. In our model, motors bind to and unbind from sites on the overlap, step toward the MT plus-end (if not sterically blocked by another motor), and switch between filaments [34, 35] (fig. 1B). Entry and exit rates characterize how motors move into/out of the overlap. MTs grow at their plus-ends in the absence of motors, but can shorten due to motor activity (fig. 1C). A steady state is reached when the average number of motors entering the overlap equals the number leaving. The steady-state overlap length therefore depends on motor density at MT plus-ends, which can be described by balancing the inward and outward fluxes at the ends [10, 11] or by a total binding constraint [34, 35] analyzed by studying the phase-space flow. Since the motor exit rate depends on microtubule shrinking, higher bulk motor concentration leads to shorter overlap length. Although previous work on microtubule length regulation found that minimal models can quantitatively agree with the data, here we find that most simple mechanisms of motor-catalyzed MT shortening give predictions at odds with the experimental data [23]. We identify one mechanism that can explain the data, motivated by recent work that has identified altered motor binding kinetics that can favor binding at MT plus-ends [36–43]. Such altered binding kinetics, along with processive depolymerization of the MT, increases the effective motor density in the overlap and changes the dependence of the steady-state overlap length on the bulk motor concentration.

We consider two antiparallel MTs with overlap length *L* (fig. 1). Each site *i* on each MT can be occupied by a motor 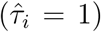 or empty 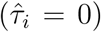. Motors in the overlap step and are sterically excluded in a totally asymmetric exclusion process (TASEP), bind and unbind according to Langmuir kinetics (LK), and switch MTs [23, 34, 35]. For simplicity, we normalize rates by the motor speed, so the stepping rate is 1. Then the mean-field dynamics of the average motor occupancy 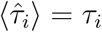 satisfy

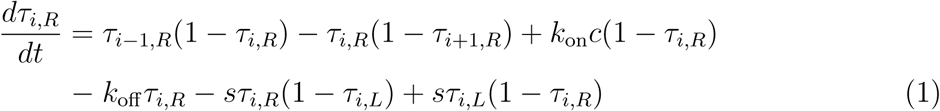

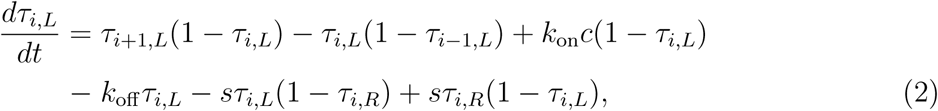

where *i* labels the site on the filament, *R, L* label the orientation of the MT, *k*_on_*c* is the normalized motor binding rate, *k*_off_ the unbinding rate, and *s* the switching rate. Parameter values were estimated from previous measurements [23] or comparison of our simulations to the data [34] (table S1 [44]).

The overlap length is dynamic and regulated by motor occupancy at the MT plus-ends [23]. In the absence of motors, MTs grow with speed *u*, but the growth speed is lowered if the plus-end site is occupied by a motor. We consider several possible models for motor activity at the plus-end. Perhaps the simplest is density-dependent shrinking [9–11, 45], which in the mean-field approximation gives an equation for MT length *L*

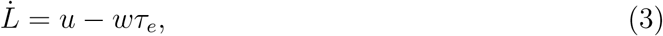

where 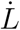 is the time derivative of the overlap length and *τ*_*e*_ is the average motor density at the plus-end. Other possible models that depend only on the end occupancy include density-dependent growth inhibition 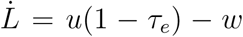, and density-dependent shrinking and growth inhibition 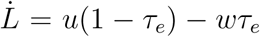. However, all of these models lead to a scaling of steady-state overlap length with bulk motor concentration of the form *L* ∼ *c*^−1^, which contradicts the experimental results [23].

To see this discrepancy, we analyze the linearized mean-field equations in the continuum limit *τ*_*i*_ *≃ ρ*(*x*). The full solution contains approximately linearly varying regions separated by domain walls of negligible width [46], allowing us to solve the linearized equations and treat domain walls as discontinuous jumps obeying a matching condition [26]. This allows an analytic analysis of the steady-state overlap length. At steady-state the linearized equations are [35]

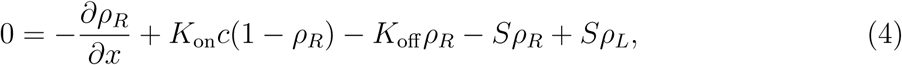

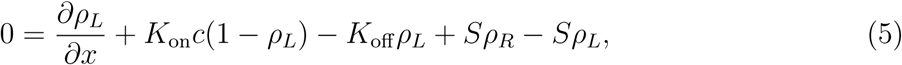

defined on an overlap of length *L* with *x* between −*L/*2 and *L/*2. The density at the plus-end of each filament is set to *ρ*, giving boundary conditions *ρ*_*R*_(−*L/*2) = 0, *ρ*_*R*_(*L/*2) = *ρ, ρ*_*L*_(−*L/*2) = *ρ, ρ*_*L*_(*L/*2) = 0 [47]. The binding/unbinding and switching rates are then 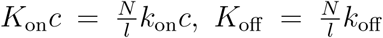, and 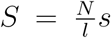, where *N* is the total number of sites in a reference length *l*. We set the reference length *l* to be the length of a physical size of a tubulin dimer, the minimal subunit of a microtubule. We define the total and difference densities *ρ*_Σ_ = *ρ*_*R*_ + *ρ*_*L*_ and *ρ*_Δ_ = *ρ*_*R*_ − *ρ*_*L*_, which and find the general solution to the linear equation *ρ*_Σ_ = *A* cosh(λ*x*)+ 2*ρ*_0_, where 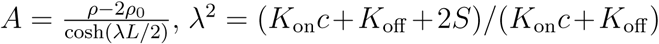, and *ρ*_0_ = *K*_on_*c/*(*K*_on_*c* + *K*_off_). The boundary density *ρ* is determined by motor flux balance at site *e* [10]:

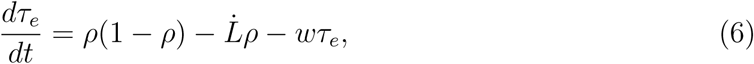

where 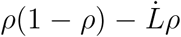 is the flux from the bulk, and *wτ*_*e*_ is the flux due to MT shortening. The system has a total binding constraint [35] (figure S1), which comes from balancing net motor entrance into and exit from the overlap. For the continuum equations, this constraint is 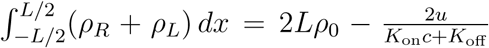, where we have assumed the density-dependent shrinking model of equation (3) at steady state, so the rate of motor exit from the overlap due to filament shortening is *wτ*_*e*_ = *u* [44]. Then the steady-state overlap length is determined by the bulk motor concentration

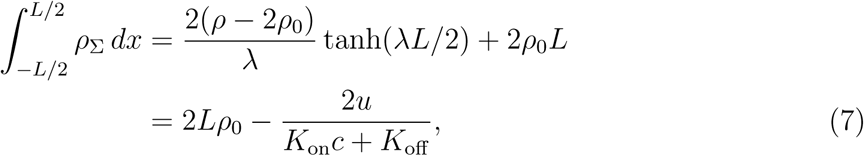

which can be inverted to relate the overlap length to the model parameters

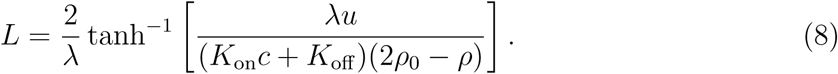

The scaling of the steady-state overlap length with bulk motor concentration can be derived through a Taylor expansion

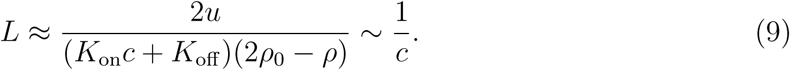

The scaling *L* ∼ *c*^−1^ differs significantly from the experimental results [23] (fig. 2). All three simple models of the end dynamics discussed above have this same scaling, suggesting that they are not sufficient to explain the data.

**FIG. 2.**
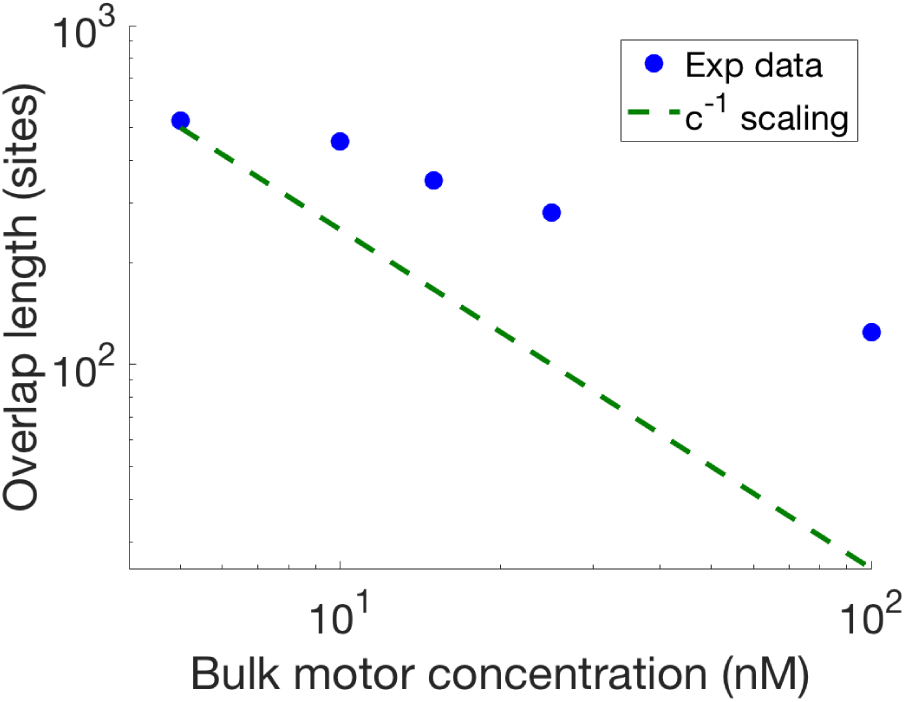
Comparison of the experimental data [23] with the *L* ∼ *c*^−1^ scaling predicted by simple length regulation mechanisms.

This discrepancy motivated consideration of other possible mechanisms by which motors at the plus-end might shorten the MT. MT structure is altered near the plus-ends, which can be recognized by kinesin motors [36–43]. Therefore we considered site-specific motor binding and unbinding rates at the end site *e* and end tracking with MT polymerization (fig. 3). We also consider processive depolymerization, controlled by the MT shrinking rate and the occupancy of the site adjacent to the end, which we label site *e*− 1 with occupancy *τ*_*e*−1_. In this model, if site *e* − 1 is occupied, the motor at end site unbinds as the MT shortens, while if site *e* − 1 is unoccupied, the end motor moves backward to the previous site as the MT shortens (fig. 3). In addition, MT length dynamics are controlled by site *e* − 1 adjacent to the end. *i*.*e*., 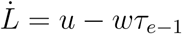. The mean-field time evolution of *τ*_*e*_ is then

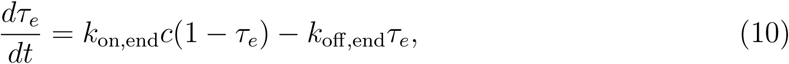

where *c* is the bulk motor concentration, and *k*_on,end_ and *k*_off,end_ are the on-rate constant and off-rate. At steady state, 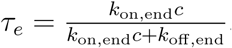. If depolymerization is processive, the site *e –* 1 mean-field occupancy is determined by

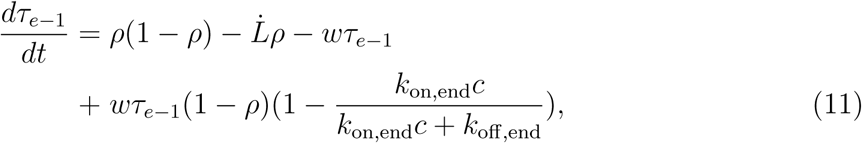

where 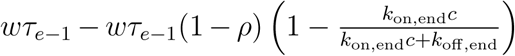 is the flux due to MT length dynamics, and the other terms are similar to those in equation (6). For small *ρ*, the steady-state outward flux becomes 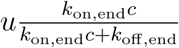 and flux balance requires that 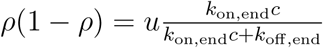.

**FIG. 3.**
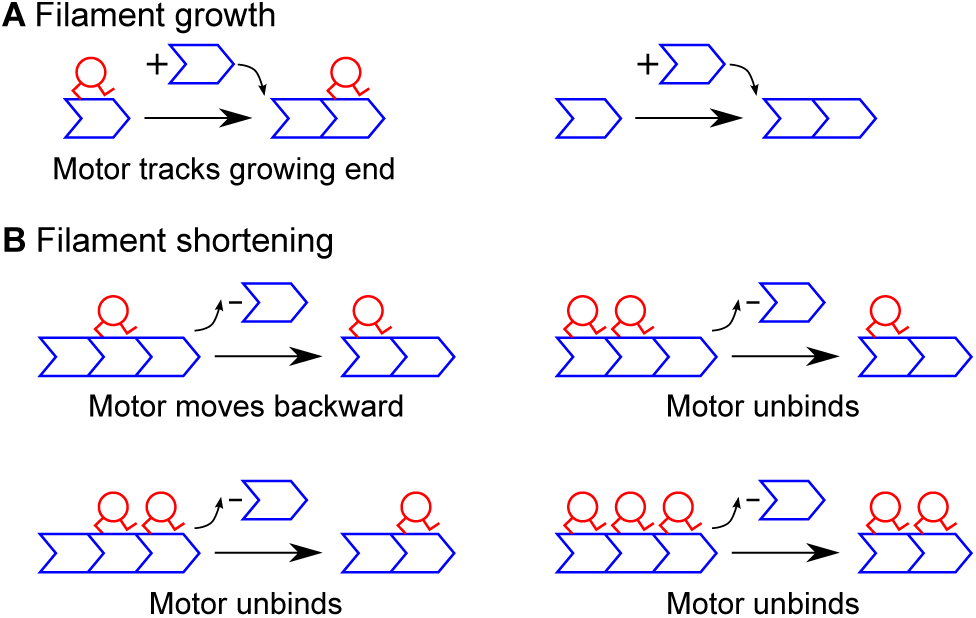
Schematic of growing and shrinking mechanisms considered in the model. (A) Filament growth. The sites shown on the left represent site *e* − 1. In this mechanism, the motor moves to a newly added site immediately. (B) Filament shrinking. The sites shown are the last site *e* and the adjacent, second-to-last site *e* − 1. Processive depolymerization occurs when site *e* − 1 is occupied but the site behind it is empty, and the motor moves backward as the final site is removed.

Using this linearized mean-field model with small-boundary density to determine the steady-state overlap length, we find that the total binding constraint is 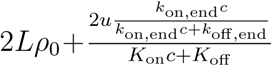 [48]. We then determine the steady-state overlap length (fig. 4), either in the linearized model (dashed green curve) or in the full nonlinear model using phase-space flow to numerically connect the boundary conditions [35] (black curve, see fig. 5). The blue points show the experimentally measured overlap lengths [23] and the red points results of kinetic Monte Carlo simulations. This version of the model shows excellent agreement with the experimental data.

**FIG. 4.**
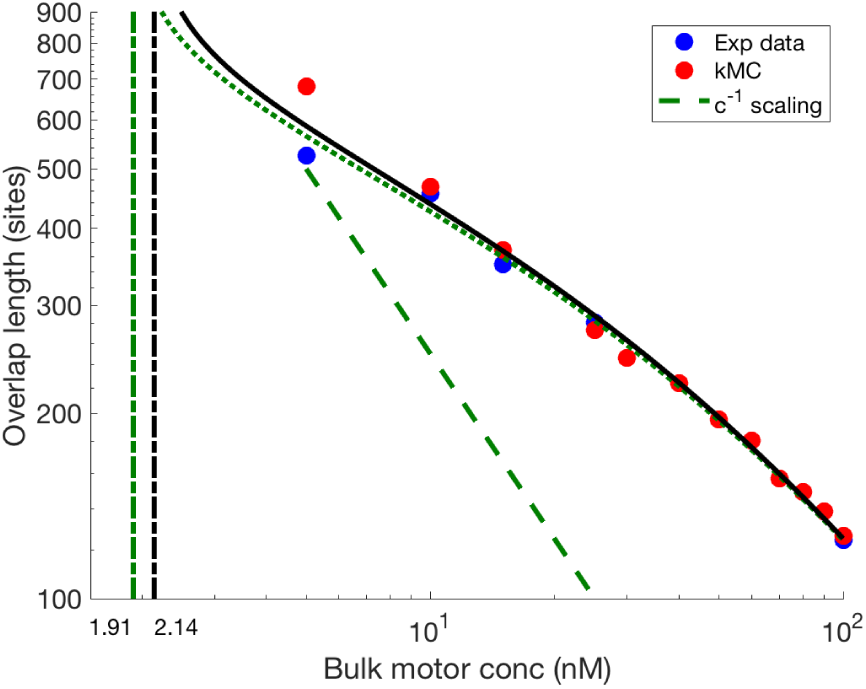
The steady-state overlap length as a function of bulk motor concentration. The green-dashed curve is from the linear model, and the black curve is the theoretical prediction using phase space flow[35] (see the main text for the detail description). The critical concentration of the black curve is around 2.2 nM. The shrinking rate is 50 *µ*m min^−1^. Other parameters are the reference values of table S1.

**FIG. 5.**
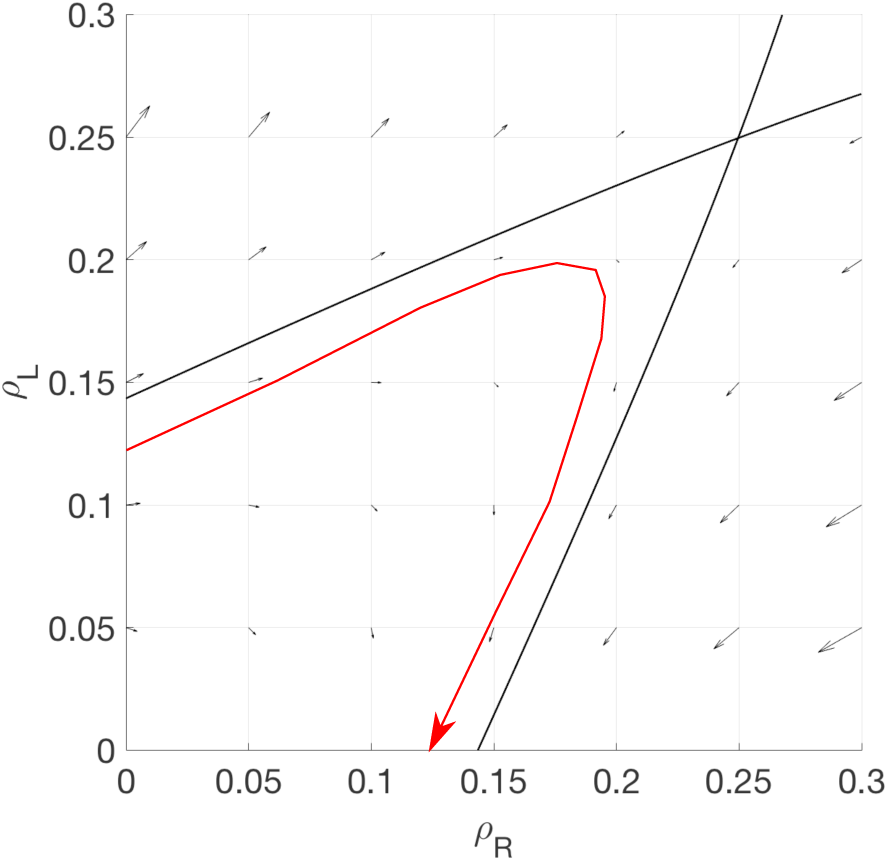
Illustration of steady-state overlap length computation using the phase space flow. Black arrows indicate the local phase space flow in the *ρ*_*R*_ − *ρ*_*L*_ plane; the black line is the flow line that crosses the Langmuir isotherm. The red curve is the path from (*ρ*_*R*_(−*L/*2), *ρ*_*L*_(−*L/*2)) = (0.125, 0) to (*ρ*_*R*_(*L/*2), *ρ*_*L*_(*L/*2)) = (0, 0.125); the integrated length of this curve is the steady-state overlap length [35].

The strong agreement between the simulations and mean-field model may be surprising, given previous work: the variable length of the overlap creates an effective boundary condition on the bulk overlap, where the balance between the boundary and the motor flux from the bulk creates a domain wall. This is characteristic of TASEP models [24–27]. Previous work has argued that the strong correlation between two nearby sites around the domain wall leads to a failure of the mean-field description [24, 25, 27]. However, in our model the binding and unbinding rates at the end are relatively large (compared to the bulk binding kinetics) at site *e*, and therefore the correlation between *τ*_*e*−1_ and *τ*_*e*_ is significant only when the shrinking rate is high. In addition, the motor density *ρ*(*L*) at the other side of the domain wall is usually small (especially when the shrinking rate is large), so the mean-field description is a good approximation.

Our theory predicts that a steady-state overlap length cannot occur if the motor density is too small. The phase-space method gives a natural interpretation of the minimum motor concentration required for a steady-state overlap: when the boundary condition passes the Langmuir isotherm point, the overlap length required to satisfy the boundary conditions becomes infinite. Similarly, in the linearized approximation the overlap length diverges at the critical concentration. The calculated critical concentration is approximately 2 nM, comparable to the result found experimentally [23].

Previous work has established the Extremal Current Principle to determine motor-density profiles in the limit that binding and unbinding can be neglected (conserved particle number) at domain walls [10, 30, 49, 50]. The ECP allows identification of phase boundaries between the low-density, high-density, and maximum-current phases of the overlap model (see Supplementary material). This analysis shows that a steady-state overlap length occurs only in the low-density phase and for a restricted range of possible MT growth speeds. Since steady-state overlaps occur for the IN phase, the domain wall is near the end of the overlap. This implies that the length between the two domain walls (one at each end) is approximately equal to the overlap length, which makes the length predicted in equation nearly equal to the overlap length. In the case of the purely density-controlled model (with no effects of the site adjacent to the end), *u* is independent of motor concentration and therefore *ρ* is independent of the bulk motor concentration. However, a motor-dependent *u* will alter the scaling of *ρ* with the bulk motor concentration. Dependence on the occupancy of the site adjacent to the MT end as we propose is one way this could occur.

In this article, we studied the length regulation of MT antiparallel overlaps in a model with motor motion and steric exclusion (TASEP), binding kinetics (LK), and switching between filaments. We used both mean-field continuum equations and kinetic Monte Carlo simulations to solve the model. Extension of the model to include plus-end-specific binding kinetics and processive depolymerization allows both the simulation and mean-field solutions agree with the experimental results [23] (fig. 4).

## ACKNOWLEDGEMENTS

We thank Matthew Glaser, Loren Hough, and Radhika Subramanian for useful discussions. This work was supported by NSF Grants No. DMR-0847685, No. DMR-1725065, and No. DMR-1551095 and NIH Grant No. K25GM110486 to M.D.B., a fellowship to H.-S.K. provided by matching funds from the NIH/CU Biophysics Training Program, and facilities of the Soft Materials Research Center under NSF MRSEC Grant No. DMR-1420736, and the Max Planck Institute for the Physics of Complex Systems for providing computing resources.

