## Supplemental text and figures for "Theory of microtubule length regulation in antiparallel overlaps"

### Supplementary material

#### I. KINETIC MONTE CARLO SIMULATION METHODS

We performed kinetic Monte Carlo (kMC) simulation of the discrete model. Motor motion in an overlap with $N$ sites is simulated with these rules:

1. Randomly choose a lane ($R/L$) and site $i$.

2. If the site is empty, attach a motor with probability $k_{\text{on}}c\Delta t$ ($k_{on,end}c\Delta t$ for site $N$ on lane $R$ and site 1 on lane $L$). If the site is occupied, detach the motor with probability $k_{\text{off}}\Delta t$ ($k_{off,end}\Delta t$ for site $N$ on lane $R$ and site 1 on lane $L$).

3. If the site is occupied and the adjacent site toward the plus end is empty, move the motor forward with probability $v\Delta t$; however, moves from site 1 or to site $N$ are forbidden.

4. If the bulk site (i.e., not site 1 or $N$) is occupied and the corresponding site on the neighboring lane is empty, switch the motor to the other lane with probability $s\Delta t$.

5. If the site is a boundary site (i.e., site 1 or $N$), it has probability $\alpha\Delta t$ (for site 1 on lane $R$ and site $N$ on lane $L$) to attach a motor, and probability $\beta\Delta t$ (site $N$ on lane $R$ and site 1 on lane $L$) to detach a motor. In this paper, $\alpha = \beta = 0$ indicate no motor coming in and out through the boundary fluxes.

6. Repeat steps 1-5 $2N$ times total to sample all sites on both lanes.

Microtubule length dynamics and regulation follow the rules:

1. Randomly choose a lane ($R/L$).

2. If second site from the plus end (i.e., site $N-1$ on lane $R$ and site 2 on lane $L$) of the lane has a motor, reduce the overlap length by 1 site with probability $w\Delta t$. If shortening does not occur, increase the overlap length by 1 site with probability $u\Delta t$.

3. Move the motor at the last site or penultimate site if necessary (see the main text fig. 3).

4. Repeat steps 1-3 2 times total to sample both plus ends.

We chose the time step $\Delta t$ to give a characteristic time for motor binding/unbinding of about $10^5$ time steps. For the reference parameter set and a bulk motor concentration of 100 nM, we used $\Delta t = 5 \times 10^{-4}$ s. Approximately $4 \times 10^7$ time steps were used to reach steady state without any microtubule growth or shrinking, and then growth and shrinking were simulated for $2 \times 10^7$ time steps. Data for figures was collected in the last $10^7$ time steps. The initial overlap length is 400 sites, and the steady-state length is typically reached within $10^7$ time steps.

To determine the average motor concentration and overlap length, we averaged $5 \times 10^4$ samples separated by 200 time steps. The reference parameter set was obtained from Bieling *et al.* [1] or estimated based on comparison of our kMC simulations to the B data [2, 3]; see table S1.

#### II. BOUNDARY CONDITIONS AND THE OTHER POSSIBLE MODELS

Here we connect the boundary conditions in the reaction-diffusion equation to the flux-balance boundary conditions of the TASEP. The steady-state continuum mean-field equations of an overlap (see equations (1), (2), (5) and (6) in the main text) are:

$$0 = -\partial_x \left(\rho_R(1-\rho_R)\right) + K_{\text{on}}c(1-\rho_R) - K_{\text{off}}\rho_R - S\rho_R + S\rho_L \tag{S1}$$
$$0 = \partial_x \left(\rho_L(1-\rho_L)\right) + K_{\text{on}}c(1-\rho_L) - K_{\text{off}}\rho_L + S\rho_R - S\rho_L, \tag{S2}$$

and the corresponding linearized continuum mean-field equation is a steady-state reaction-diffusion equation:

$$\frac{\partial^2}{\partial x^2}(\rho_R+\rho_L) = \frac{\partial^2 \rho_\Sigma}{\partial x^2} = (K_{\text{on}}c + K_{\text{off}} + 2S)(K_{\text{on}}c + K_{\text{off}})\rho_{tot} - 2K_{\text{on}}c(K_{\text{on}}c + K_{\text{off}} + 2S). \tag{S3}$$

| Symbol | Parameter | Reference value | Notes |
|---|---|---|---|
| $v$ | Motor speed | 0.5 $\mu$m s$^{-1}$ | Measured by Bieling et al. [1] |
| $u$ | Microtubule growing rate | 2.4 $\mu$m min$^{-1}$ | Estimated based on the growing rate without motors [1] |
| $w$ | Microtubule shrinking rate | 50 $\mu$m min$^{-1}$ | |
| $K_{\rm on}$ | Binding rate constant | $2.811 \times 10^{-4}$ nM$^{-1}$ s$^{-1}$ | Estimated based on motor density profiles and kymographs in Kuan and Betterton [2] |
| $c$ | Bulk motor concentration | 1–100 nM | Varied by Bieling et al. [1] |
| $K_{\rm off}$ | Unbinding rate | 0.169 s$^{-1}$ | Measured by Bieling et al. [1] |
| $k_{on,end}$ | Binding rate at the $\tau_{cap}$ site (see main text) | $v/10$ | Estimated based on our model and the length regulation results from Bieling et al. [1] paper. |
| $k_{off,end}$ | Binding rate at the $\tau_{cap}$ site (see main text) | $82 \times v/10$ | Estimated based on our model and the length regulation results from Bieling et al. [1] paper. |
| $s$ | Switching rate | 0.44 s$^{-1}$ | Measured by Bieling et al. [1] |
| $\alpha$ | Motor flux constant into overlap from MT minus end | 0 | Motors bind primarily inside the overlap; see discussion in Kuan and Betterton [2]. We varied $\alpha$ between 0 and 1 to determine the model phase diagram |
| $\beta$ | Motor flux constant out of overlap from MT plus end | 0 | An upper bound on the end motor unbinding rate is $\beta = 2.811 \times 10^{-4}$; see Kuan and Betterton [2]. We varied $\beta$ between 0 and 1 to determine the model phase diagram |

TABLE S1. Parameter values for the reference parameter set, taken from experimental measurements or estimated as noted.

The general solution is $\rho_\Sigma = A\cosh(\lambda x) + 2\rho_0$, assuming symmetry in the overlap $\rho_\Sigma(x) = \rho_\Sigma(-x)$ (which elimiates the hyperbolic sine term). Here $\lambda^2 = (K_{\rm on}c + K_{\rm off} + 2S)(K_{\rm on}c + K_{\rm off})$.

Because the linearized equation is derived from the TASEP, one might expect that any inconsistency between these two descriptions comes from linearization. However, linearization effects can be minimized by taking advantage of the appearance of domain walls in the TASEP: we write the solution to the linearized equation in pieces separated by the domain walls [4]. Each piece of the solution is typically well-described by the linearized equation. (We note that any disagreement can be addressed by phase-space flow analysis [3].) The domain walls can then be treated as discontinuous jumps that satisfy a matching condition [4].

The total binding constraint [3] connects the linearized equation and the boundary condition of the TASEP gives the length regulation. For the total binding constraint, the overall motor density for a given overlap length $L$ is derived by integrating equations (S1) and (S2):

$$\int_{-L/2}^{L/2} \rho_\Sigma \, dx = 2L\rho_0 - \frac{2C}{K_{\rm on}c + K_{\rm off}}, \tag{S4}$$

where $\rho_0 = K_{\rm on}c/(K_{\rm on}c + K_{\rm off})$ is the steady-state motor density due to the binding kinetics (LK), and $-\frac{2C}{K_{\rm on}c+K_{\rm off}}$ is the density change due to the boundary condition. The constant $C$ can be understood as the boundary flux, which is clearer if written as $C = \rho(1-\rho)$. If the domain wall is at the boundary (or, for the trivial case, there is no domain wall), the overall motor density through the linearized equation links the undetermined constant $A$ in the general solution to the flux $C$:

$$\int_{-L/2}^{L/2} \rho_\Sigma \, dx = \frac{2A\sinh[\frac{\lambda L}{2}]}{\lambda} + 2L\rho_0$$

$$\Rightarrow A = -\frac{C\lambda}{\sinh[\frac{\lambda L}{2}]\,(K_{\rm on}c + K_{\rm off})}. \tag{S5}$$

If the boundary condition of the linearized equation is $\rho_\Sigma(L/2) = \rho$, then $A$ becomes $\frac{\rho - 2\rho_0}{\cosh[\lambda L/2]}$ for the general solution and the length as the function of the boundary flux $C$ and bulk motor concentration $c$ is

$$L = \frac{2}{\lambda}\tanh^{-1}\left[\frac{\lambda C}{(2\rho_0 - \rho)\,(K_{\rm on}c + K_{\rm off})}\right]. \tag{S6}$$

Besides the normal TASEP boundary condition ($\alpha$ and $\beta$), the length regulation can induced the effective boundary condition. The general length regulation equation is

$$\frac{\partial L}{\partial t} = \bar{u}(\{\tau_i\}) - \bar{w}(\{\tau_i\}), \tag{S7}$$

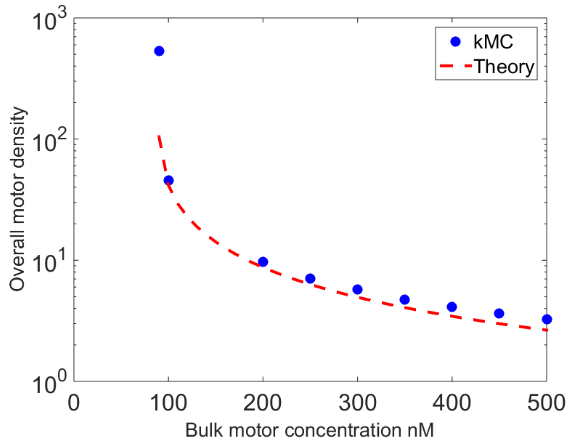


FIG. S1. Total binding as a function of bulk motor concentration. The growing rate is 2 $\mu$m min$^{-1}$and the shrinking rate is 40 $\mu$m min$^{-1}$. Other parameters are the reference values of table S1.

where $\bar{u}$ is the growing speed, $\bar{w}$ is the shrinking speed, and $\{\tau_i\}$ indicate they can be functions of motor density at any site(s). At steady state, $\bar{u} = \bar{w}$.

##### A. Length regulation by the end-site density

The dependence of $\bar{u}$ and $\bar{w}$ on the motor end occupancy could occur in several forms:

1. Density-controlled shortening $\frac{\partial L}{\partial t} = u - w\tau_e$,
2. Density-controlled growth inhibition $\frac{\partial L}{\partial t} = u(1-\tau_e) - w$,
3. Density-controlled shortening and growth inhibition $\frac{\partial L}{\partial t} = u(1-\tau_e) - w\tau_e$.

In these models, the steady-state solutions are (1) $\tau_e = \frac{u}{w}$, (2) $\tau_e = \frac{u-w}{w}$, and (3) $\tau_e = \frac{u}{u+w}$. The motor exit rate is then a constant, $w\tau_e$ [5], which leads to a constant $\rho$ in the linearized equation and $C = w\tau_e$.

We note that the motor exit rate $w\tau_e$ is the flux of the linearized equation at the boundary $\frac{\partial \rho_{tot}}{\partial x}\Big|_{x=L/2} = \lambda A \sinh[\frac{\lambda L}{2}] = -\frac{C\lambda^2}{K_{\rm on}c+K_{\rm off}} = \frac{-w\tau_e}{K_{\rm on}c+K_{\rm off}+2S} \sim -\frac{w\tau_e}{2S}$, where the last approximation is due to the relative large switching process rather than the binding/unbinding processes.

The scaling behavior of the steady-state overlap length as a function of bulk motor concentration can be derived by Taylor expansion, because the motor stepping rate is much larger than the rates of motor binding/unbinding and switching. Therefore

$$L = \frac{2}{\lambda}\tanh^{-1}\left[\frac{\lambda w\tau_e}{(K_{\rm on}c + K_{\rm off})(2\rho_0 - \rho)}\right] \approx \frac{2w\tau_e}{(K_{\rm on}c + K_{\rm off})(2\rho_0 - \rho)} \sim \frac{1}{c}, \tag{S8}$$

which implies the all three density-controlled models give the same scaling.

##### B. Length regulation by the two-point density

Rather than alterations to MT dynamics determined by the motor density at one site at the MT end, dynamics could be controlled by the two-point densit. One possible two-point model is

$$\frac{\partial L}{\partial t} = u - w\langle\tau_N\tau_{N-1}\rangle, \tag{S9}$$

where the subscripts $N$ and $N-1$ denote the last and second-to-last site. The steady-state solution gives $w\langle\tau_N\tau_{N-1}\rangle = u$, and therefore a constant $\rho$. As a result, the steady-state length has the same scaling behavior as for the single-site models.

##### C. Length regulation by multiple protofilaments

The models considered so far represent the MT by a single protofilament. In principle coupling between multiple protofilaments could alter the dependence of MT dynamics of motor occupancy. If we consider a two-protofilament MT, one length-regulation model is

$$\frac{\partial L}{\partial t} = u - w\tau_{e,1}(1-\tau_{e,2}) - w\tau_{e,2}(1-\tau_{e,1}) - w'\tau_{e,1}\tau_{e,2}, \tag{S10}$$

where $\tau_{e,i}$ is the motor density at the end at protofilament $i$. If the two protofilaments are physically and chemically the same, there are only two possible configurations that can lead to shortening: $(\tau_{e,1}, \tau_{e,2}) = (1,0)$ or $(0,1)$ and $(\tau_{e,1}, \tau_{e,2}) = (1,1)$. In principle each of these cases could have a different MT shortening rate. The motor exit rate from protofilament 1 is

$$[w\tau_{e,1}(1-\tau_{e,2}) + w\tau_{e,2}(1-\tau_{e,1}) + w'\tau_{e,1}\tau_{e,2}]\,\tau_{e,1} = w\tau_{e,1}(1-\tau_{e,2}) + w'\tau_{e,1}\tau_{e,2}, \tag{S11}$$

where $\tau_{e,1}^2$ terms are absent because there is no difference between $\tau_{e,1}$ and $\tau_{e,1}^2$ in the microscopic model. However, the motor exit rate is still equal to a constant $u$ at steady state, leading to the same scaling with bulk motor concentration for the other models discussed.

#### III. THE EX, IN, AND MC PHASES

The motor density profile in the overlap can be determined by the Extremal Current Principle (ECP) [6–9] if motor binding/unbinding kinetics are neglected at (around) the domain wall. Using the ECP, the collective velocity $v_{collect} = \partial_\rho J$ determines the local stability of the density profile, and the shock velocity $v_{shock} = \frac{J_{left} - J_{right}}{\rho_{left} - \rho_{right}}$ determines the domain wall direction.

In our system, steady-state solution of equation (13) (in the main text) leads to $\tau_{e-1} = \frac{\rho(1-\rho-u)}{w\tau_e(1-\rho)}$. In general, ones can expect three phases: IN, EX, and MC, which relates to the three phases in the general TASEP models [6, 10]. The IN phase refers to the motor density depends on the minus end, and the EX phase is the opposite. The MC phase refers to the maximum current phase. Our fluxes at the site $e-1$ are $J_{left} = \rho(1-\rho) + \dot{L}\rho$ and $J_{right} = w\tau_{e-1} - w\tau_{e-1}(1-\rho)(1-\tau_e)$, and $\tau_{e-1}$ as the function of $\rho$ is related to the phases of it. For the MC phase, the phase due to the motor leaving flux $J_{right}$ is when $\partial_\rho J_{right} = 0$:

$$\partial_\rho J_{right} = \partial_\rho(w\tau_{e-1} - w\tau_{e-1}(1-\tau_e)(1-\rho)) = \partial_\rho\left(\frac{\rho(1-\rho-u)(\tau_e+\rho-\tau_e\rho)}{\tau_e(1-\rho)}\right) \tag{S12}$$

$$\Rightarrow \partial_\rho J_{right} = \frac{(1-\rho_{MC})^2(2\rho_{MC} - \tau_e(u+2\rho_{MC}-1)+u) - u}{\tau_e(1-\rho_{MC})^2} = 0 \tag{S13}$$

$$\Rightarrow (1-\rho_{MC})^2(2\rho_{MC} - \tau_e(u+2\rho_{MC}-1)+u) = u, \tag{S14}$$

and the solution of equation (S14) leads to a $\tau_{e-1}$ in the MC phase.

In the EX phase, the right boundary condition dominates the density profile, so $\tau_{e-1} = \rho$. Therefore

$$\tau_{e-1} = \frac{\tau_{e-1}(1-\tau_{e-1}-u)}{w\tau_e(1-\tau_{e-1})} \Rightarrow \tau_{e-1} = 1 - \frac{u}{1-w\tau_e}. \tag{S15}$$

The phase boundaries between these three phases could be determined by the shock velocity. The phase boundary between IN–EX is

$$v_{shock} = \frac{\rho(1-\rho) - u\rho + w(1-\frac{u}{1-w\tau_e})\rho - w(1-\frac{u}{1-w\tau_e}) + w(1-\frac{u}{1-w\tau_e})(1-\tau_e)(1-\rho)}{\rho - (1-\frac{u}{1-w\tau_e})} = 0 \tag{S16}$$

$$\Rightarrow \rho = \tau_e w, \tag{S17}$$

which indicates the only possibility for an EX phase at the site $e-1$ is the constant bulk density $\tau_e w$ and is not possible to fulfill under different bulk motor density $c$ and MT length. The phase boundary between the IN and the MC phase is determined by (S14), and the boundary between the EX and the MC phase is the same as equation (S16) with $\rho$ replaced by $\rho_{MC}$, the density of the maximum current phase. Therefore, not all phases can exist for a steady-state MT. In particular, the EC and MC phases cannot support a constant-length overlap because of the lack

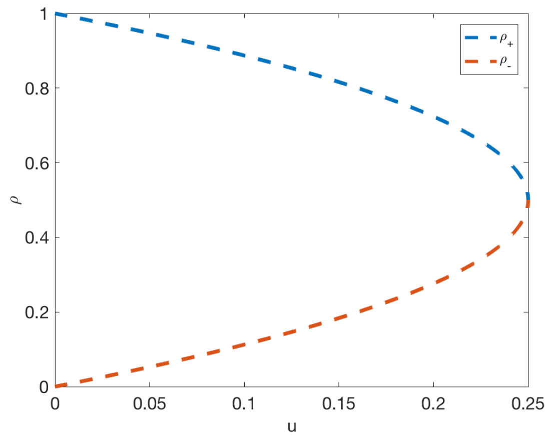


FIG. S2. Two solutions in equation (S18), $\rho_+$ is the plus solution and $\rho_-$ is the minus solution. $\rho_-$ is the only stable solution. We note that since the domain wall width approaches zero but is finite, we choose the value of motor density near the end, 5 sites from the ends.

of variation of MT plus-end density with MT length, and only in the IN phase is can the MT plus-end density vary with overlap length, allowing a nonzero steady-state length.

In the IN phase, $\tau_{e-1}$ is always $u/w$ at steady state, and $\rho$ is given through equation (8) (see the main text). This makes the IN phase the relevant one for comparison to experiment. If there depolymerization is unprocessive ($\tau_{e-1} = 1$), we find

$$\rho = \frac{1 \pm \sqrt{1-4u}}{2}. \tag{S18}$$

This has two solutions, and $\rho_-$ is the only stable solution (Figure S2). To see that the $\rho_-$ is unstable, note that since $\rho_+ > 0.5$, its corresponding phase-space velocity [3] is smaller if the motor density is higher while the flux is lower. The overlap length is longer if the motor density at the ends is higher, and vice versa. If the motor density at ends is higher than $\rho_+$, the flux decreases, which makes the end motor density lower, and thus the overlap becomes longer. If the motor density at ends is lower than $\rho_+$, the flux increases, and makes the overlap shrink. Thus, $\rho_+$ is an unstable solution. Conversely, for the $\rho_-$ solution, the overlap length is shorter if the motor density at ends is higher, and it is longer if the motor density at ends is smaller. Since the flux increases if the motor density is greater than $\rho_-$ or decreases if the motor density is smaller than $\rho_-$, the overlap length will return to $\rho_-$.

[1] P. Bieling, I. A. Telley, and T. Surrey, Cell **142**, 420 (2010).
[2] H.-S. Kuan and M. D. Betterton, Biophysical Journal **110**, 2034 (2016).
[3] H.-S. Kuan and M. D. Betterton, Phys. Rev. E **94**, 022419 (2016).
[4] A. Parmeggiani, T. Franosch, and E. Frey, Phys. Rev. E **70**, 046101 (2004).
[5] The motor exit rate is not $w\tau_e^2$ but $w\tau_e$ in case (1) and (3) is because there is no difference between $\tau_e$ and $\tau_e^2$ in the microscopic model. In other words, microscopically, $\tau_e$ and $\tau_e^2$ both behave like $\tau_e$ since $\tau_e$ can only be 0 or 1.
[6] J. Krug, Phys. Rev. Lett. **67**, 1882 (1991).
[7] A. B. Kolomeisky, G. M. Schütz, E. B. Kolomeisky, and J. P. Straley, J. Phys. A: Math. Gen. **31**, 6911 (1998).
[8] V. Popkov and G. M. Schütz, Europhys. Lett. **48**, 257 (1999).
[9] A. Melbinger, L. Reese, and E. Frey, Phys. Rev. Lett. **108**, 258104 (2012).
[10] B. Derrida, E. Domany, and D. Mukamel, Journal of Statistical Physics **69**, 667 (1992).
